# Variable orthogonality of RDF – large serine integrase interactions within the ϕC31 family

**DOI:** 10.1101/2024.04.03.587898

**Authors:** Alasdair I. MacDonald, Aron Baksh, Alex Holland, Heewhan Shin, Phoebe A. Rice, W. Marshall Stark, Femi J. Olorunniji

## Abstract

Large serine integrases are phage- (or mobile element-) encoded enzymes that catalyse site-specific recombination reactions between a short DNA sequence on the phage genome (*attP*) and a corresponding host genome sequence (*attB*), thereby integrating the phage DNA into the host genome. Each integrase has its unique pair of *attP* and *attB* sites, a feature that allows them to be used as orthogonal tools for genome modification applications. In the presence of a second protein, the Recombination Directionality Factor (RDF), integrase catalyses the reverse, excisive reaction, generating new recombination sites, *attR* and *attL*. In addition to promoting *attR* x *attL* reaction, the RDF inhibits *attP* x *attB* recombination. This feature makes the directionality of integrase reactions programmable, allowing them to be useful for building synthetic biology devices. In this report, we describe the degree of orthogonality of both integrative and excisive reactions for three related integrases (ϕC31, ϕBT1, and TG1) and their RDFs. Among these, TG1 integrase is the most active, showing near complete recombination in both *attP* x *attB* and *attR* x *attL* reactions, and the most directional in the presence of its RDF. Our findings show that there is varying orthogonality among these three integrases – RDF pairs: ϕC31 integrase was the least selective, with all three RDFs activating it for *attR* x *attL* recombination. Similarly, ϕC31 RDF was the least effective among the three RDFs in promoting the excisive activities of the integrases, including its cognate ϕC31 integrase. ϕBT1 and TG1 RDFs were noticeably more effective than ϕC31 RDF at inhibiting *attP* x *attB* recombination by their respective integrases, making them more suitable for building reversible genetic switches. AlphaFold-Multimer predicts very similar structural interactions between each cognate integrase – RDF pair. The binding surface on RDF is much more conserved than the binding surface on integrase, an indication that specificity is determined more by the integrase than the RDF. Overall, the observed weak integrase/RDF orthogonality across the three enzymes emphasizes the need for identifying and characterizing more integrase – RDF pairs. Additionally, the ability of a particular integrase’s preferred reaction direction to be controlled to varying degrees by non-cognate RDFs provides a path to tunable, non-binary genetic switches.

## Introduction

Large serine integrases catalyse site-specific recombination reactions between short DNA sequences on temperate phages (*attP*) and equivalent sequences on the genome of their bacterial hosts (*attB*) (Olorunniji *et al*., 2016; Stark, 2017). The reaction results in the generation of new sequences called *attR* and *attL* flanking the prophage integrated into the host genome. In the reverse reaction, another phage-encoded protein called the recombination directionality factor (RDF) binds to the integrase and switches its specificity, allowing it to recombine *attR* and *attL*, regenerating *attP* and *attB* sites and excising the prophage DNA from the host genome (Figure 1). The RDF also inhibits further *attP* x *attB* recombination. These two reactions are part of the natural lysis/lysogeny cycle of temperate phages. The mechanism of catalysis of recombination by serine integrases follows the pathway established for the serine recombinase family, in which the two recombining DNA sites are held together by a tetramer of recombinase protein subunits in a synaptic complex (Figure 1). The recombination steps involving DNA cleavage, subunit rotation, and DNA religation occur within the synaptic complex, ensuring tight regulation of the chemical events and conformational changes involved (Rice, 2015; Smith, 2015).

**Figure 1:**
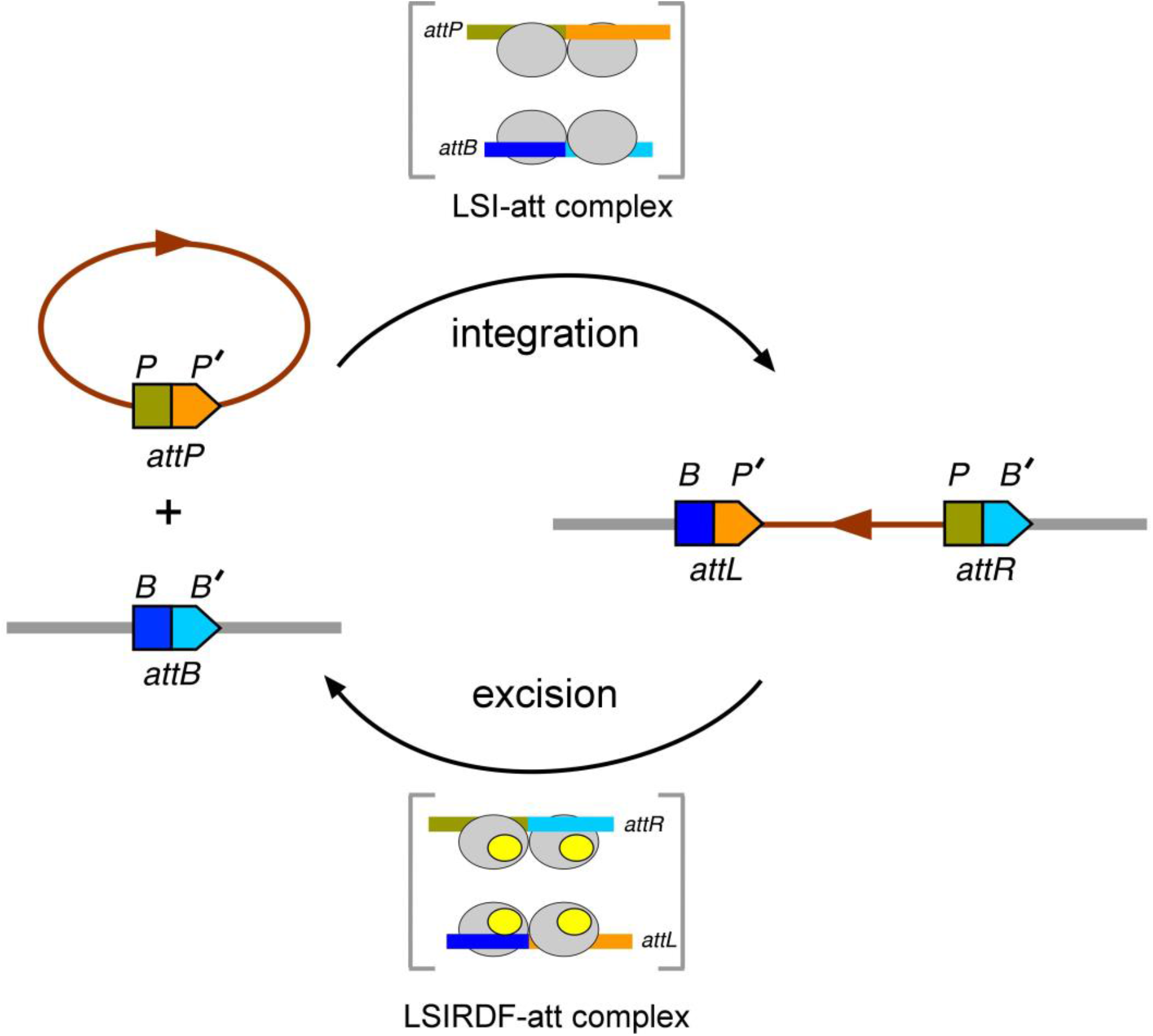
Integrative and excisive recombination reactions catalysed by large serine integrases (LSIs). LSIs integrate DNA bearing an *attP* site into a genomic location that harbours an *attB* site. In the reverse excision reaction, the recombination directionality factor (RDF) binds to the LSI and modify its specificity to catalyse *attR* x *attL* recombination. In both reactions, synapsis of *att* sites and catalysis of DNA strand exchange happen within a synaptic complex involving a tetramer of the recombinase.

Due to the unidirectional nature of the integration reaction and the abundance of serine integrases, each with its own unique sequence specificity, this group of enzymes offer the potential for developing powerful genome editing tools (Fogg *et al*., 2014; Olorunniji *et al*., 2016). Serine integrases carry out complete DNA cutting and rejoining in the recombination reaction without leaving any broken ends behind, and hence do not rely on endogenous host systems to fix broken DNA ends. Serine integrases have been adapted for a wide range of applications, including insertion of foreign DNA into specific sequences in the genomes of cells, plants and animals and the construction of genetic logic gates (Olorunniji *et al*., 2016; Stark, 2017; Colloms *et al*., 2013; Muroi *et al*., 2013; Zhang *et al*., 2021; Blanch-Asensio *et al*., 2022; Yarnall *et al*., 2023; Ginsburg *et al*., 2005; Herisse *et al*., 2018). Some tools, such as the SIRA method for assembly of replicons, require a panel of serine integrases with orthogonal sequence specificity. The ability of an RDF to trigger reversal of a particular integrative recombination renders serine integrases even more versatile as genetic tools.

ϕC31 integrase (605 amino acid residues) is the prototype serine integrase and was the first to be fully characterised *in vivo* and *in vitro* (Thorpe & Smith, 1998). It is derived from the *Streptomyces* phage ϕC31, and it is used extensively for genetic manipulations in bacteria and several eukaryotic systems. TG1 integrase and ϕBT1 integrase are two other integrases derived from *Streptomyces*; their properties and applications have been reported in the literature (Zhang *et al*., 2008; Morita *et al*., 2009a; 2009b; Du *et al*., 2015). The RDFs for all three of these integrases have been identified, and the activities for ϕC31 and ϕBT1 have been characterized both *in vivo* and in *vitro* (Khaleel *et al*., 2011; Zhang *et al*., 2013)

These three related integrases and their RDFs show significant sequence similarities, yet they have different att site sequence specificities. Hence, they are ideal candidates for analysis of the structural and biochemical basis for integrase-*att* site recognition and catalysis. They also have potential as orthogonal tools for synthetic biology applications requiring multiple serine integrases.

TG1 integrase (619 amino acid residue) is derived from the TG1 actinophage isolated from *Streptomycetes*. The integrase gene, its recombination *attB* and *attP* sites, and its activity in *E. coli* were first described by Morita *et al*. (2009a). The minimal att site requirements and *in vitro* activities were also reported by the same group (Morita *et al* 2009b).

Similarly, ϕBT1 integrase (594 amino acid residue) is derived from a phage from *Streptomyces coelicolor* (Gregory & Smith, 2003), and its activity *in vivo* and *in vitro* as well as the minimal *attB* and *attP* sequences were established by Zhang *et al*. (2008).

Zhang *et al*. (2013) found that the RDFs for ϕBT1 and ϕC31 integrases were fully exchangeable in *in vitro* recombination reactions, despite the two integrases sharing just 26% amino acid sequence identity. It was reasoned that this could be due to the 85% similarity between the two RDFs. In contrast, TG1-RDF shows 60% and 62% similarity to ϕBT1-RDF and ϕC31-RDF, respectively. To date, the effects of TG1-RDF on ϕBT1 and ϕC31 integrases have not been reported.

Since the three RDFs have regions of highly conserved sequences, they may cross-react with each other’s integrases, as has been reported for ϕC31 and ϕBT1 RDFs (Zhang *et al*., 2013). Any significant cross-reactivity across these integrases will have implications for their simultaneous (or concurrent) use in *in vivo* and *in vitro* applications. However, a full analysis of the degree of cross reactivity across these related integrase/RDF pairs have not been reported.

To clarify the extent to which the three integrases and their RDFs could be used together in synthetic biology applications, we investigated the orthogonality of the integrases and their RDFs in recombination reactions. The findings could help shed some light on the structural basis for integrase-RDF recognition and orthogonality. We also investigated whether the nature of the integrase-RDF specificities changes when the integrase and RDF are covalently joined as integrase-RDF fusions (Olorunniji *et al*., 2017). We anticipated that the findings would lay the foundation for further studies aimed at understanding the factors that determine specificity of integrase-RDF interactions.

## Results

### Sequence alignment and predicted structures of ϕC31, ϕBT1, and TG1 integrases and their recombination directionality factors (RDFs)

Sequence alignments (Figures 2 and 3) show that the three RDFs are more closely related to one another than the integrases. ϕBT1 integrase is only 24-25% identical to TG1 and ϕC31 integrases, which are 50% identical to one another. In contrast, among the RDFs, TG1 RDF is the outlier, with 61-63% identity to the other two, which are 85% identical to one another.

**Figure 2:**
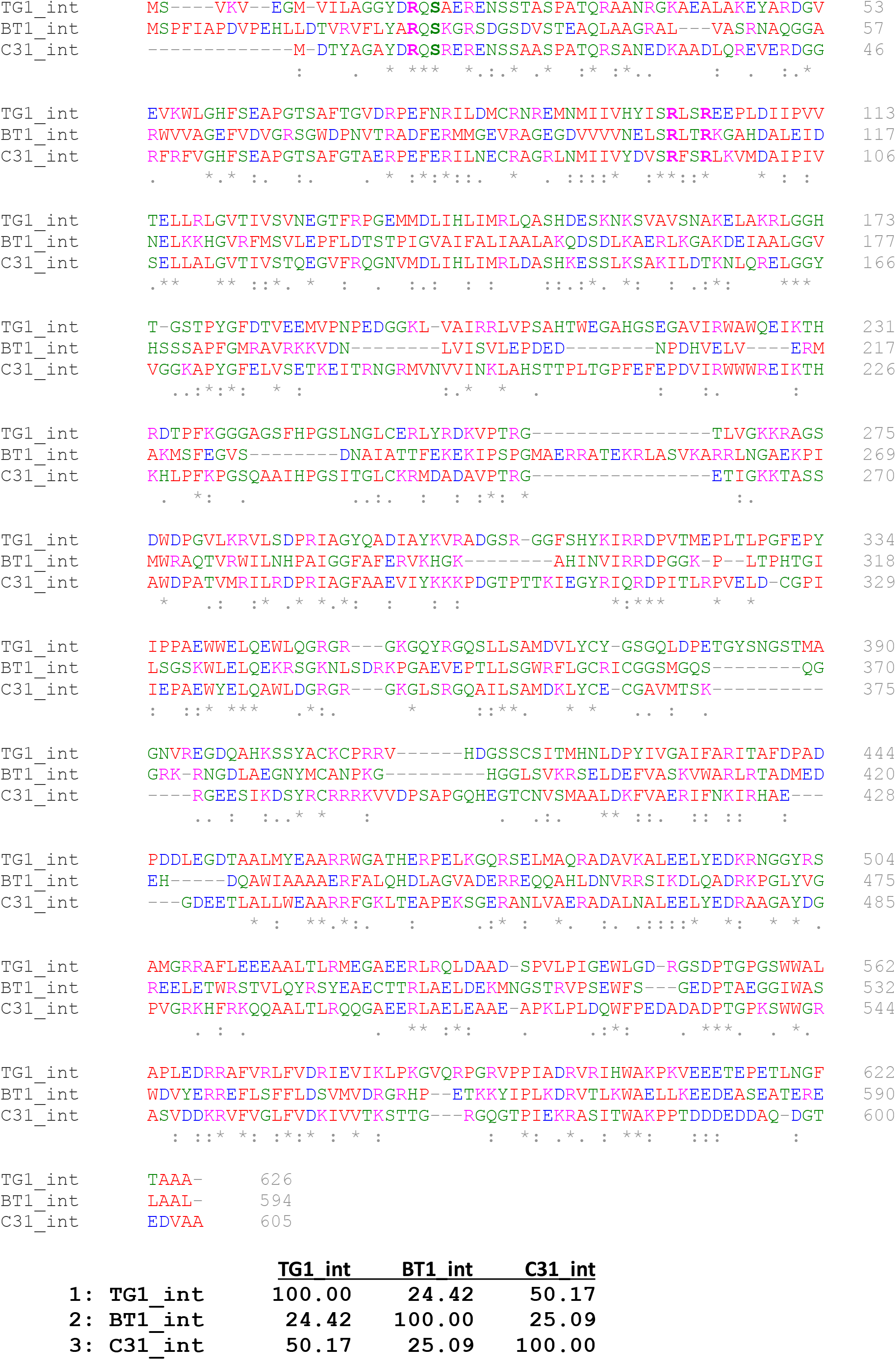
Sequence alignments of TG1, ϕBT1, and ϕC31 integrases. Key amino acid residues in the active site including the nucleophilic serine are highlighted in bold. Multiple sequence alignment was generated using Clustal Omega (Sievers et al., 2011). A matrix of pairwise percent identities is shown below. (https://www.ebi.ac.uk/jdispatcher/msa).

**Figure 3:**
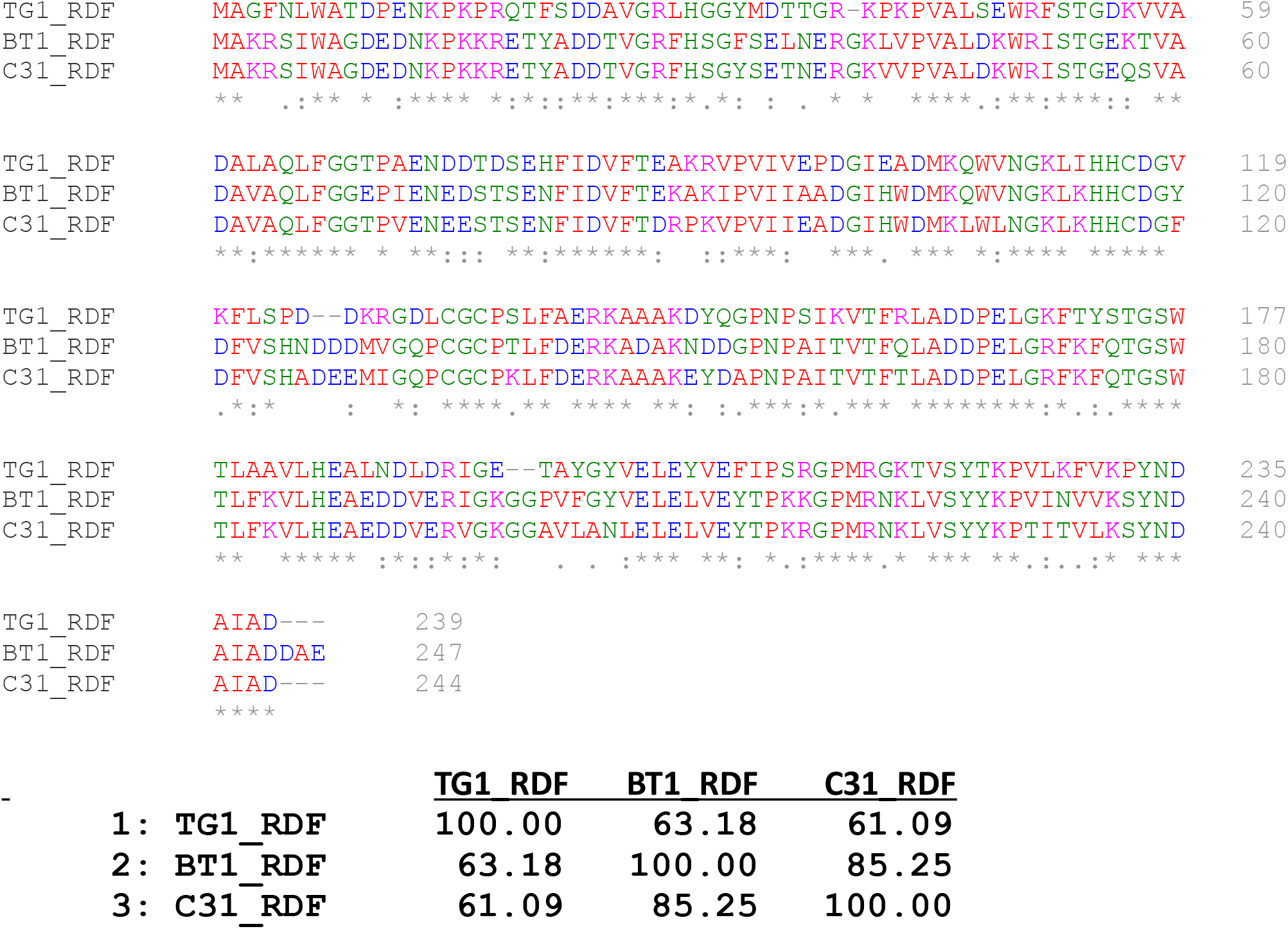
Sequence alignments of TG1, ϕBT1, and ϕC31 Recombination Directionality Factors (RDFs). Multiple sequence alignment was generated using Clustal Omega (Sievers et al., 2011). A matrix of pairwise percent identities is shown below. (https://www.ebi.ac.uk/jdispatcher/msa).

To understand the nature of integrase-RDF interactions, we used AlphaFold2-multimer to model the structures of the three integrases in complex with their cognate RDFs. As expected, the predicted structures of the two DNA binding domains are very similar to the experimental structure of A118-Int (Rutherford *et al*., 2013), which was used to model their binding to DNA. (Figure 4). All three complex models are quite similar and predict that the RDF uses a set of loops to clamp onto a hinge region between the integrases’s second DNA binding domain (DBD2; sometimes called the zinc-binding domain) and the coiled coil that is inserted within it. (Figure 4). These models are in good general agreement with prior experimental data (Mandali *et al*., 2017; Abe *et al*., 2021; Fogg *et al*., 2018).

**Figure 4:**
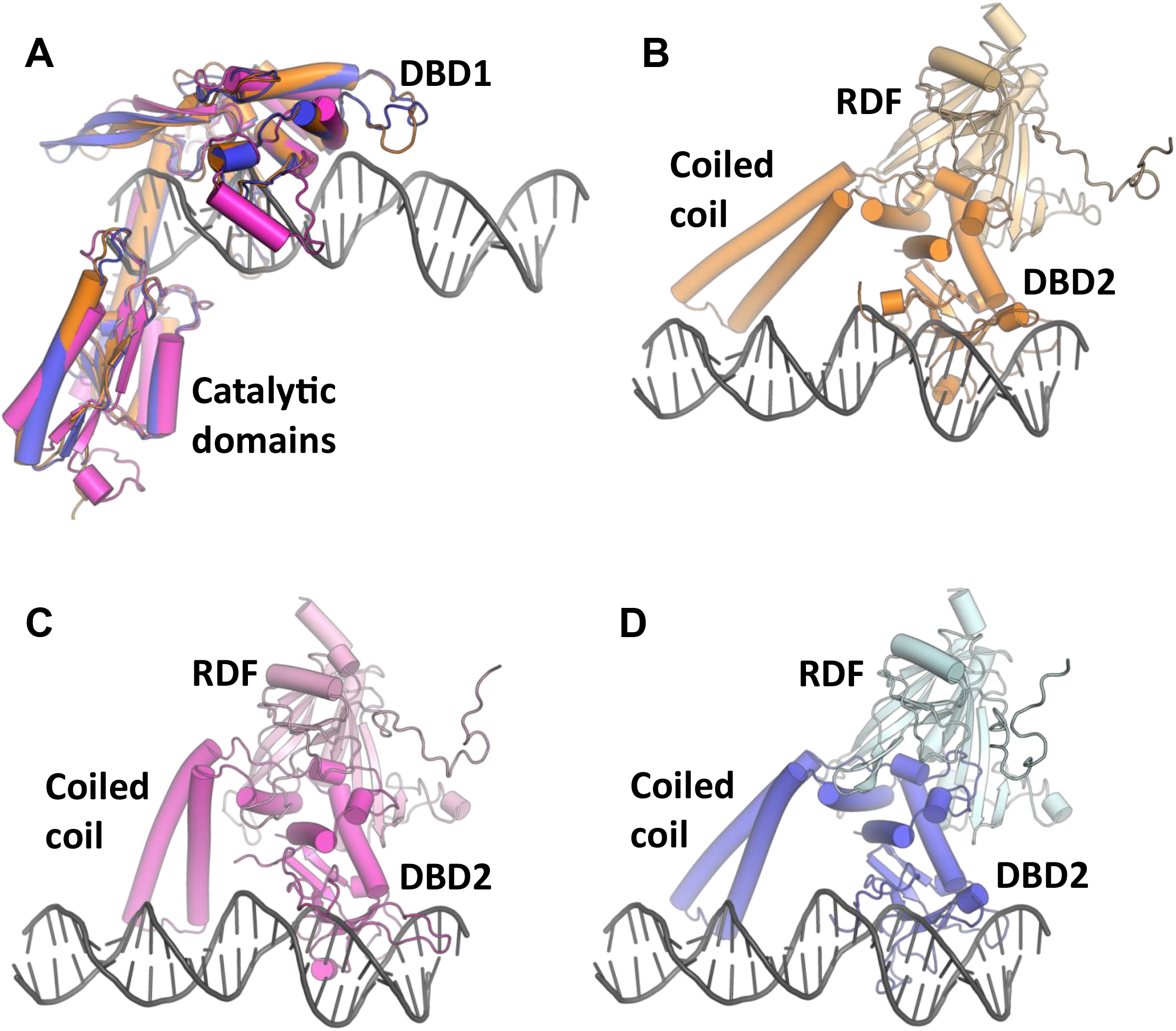
AlphaFold2-multimer structures of serine integrases and their RDFs. (**A**) AlphaFold2-predicted models for the N-terminal catalytic domains and the first DNA-binding domains (DBD1) of ϕC31 (blue), ϕBT1 (magenta), and TG1 (brown) integrases. These models were calculated using full-length protein sequences. Because the flexible linkers between domains made overall superpositions visually confusing, the models were then mapped domain-wise (except for the coiled coil) onto the individual domains of 4kis.pdb, which is a complex of the DNA binding domains of A118 integrase with half of attP. The DNA from 4kis is shown to guide the eye. (**B**-**D**). The second DNA-binding domains (DBD2) of TG1 (**B**), ϕBT1 (**C**), and ϕC31 (**D**) and their RDFs, taken from the same predicted structures as in part (A) but shown separately for clarity. Structures were generated using AlphaFold Multimer as described in Methods section. Flexible C-terminal extensions of integrases were removed for clarity.

### Recombination activities of ϕC31, ϕBT1, and TG1 integrases

*In vivo*, all three integrases catalysed *attP* x *attB* recombination to near completion, and as expected did not act on the *attR* x *attL* substrate in the absence of the RDF (Figure 5), showing the strict directionality as well as efficiency of the integration reactions. A similar pattern was observed *in vitro*, with recombination being generally efficient (Figure 6).

**Figure 5:**
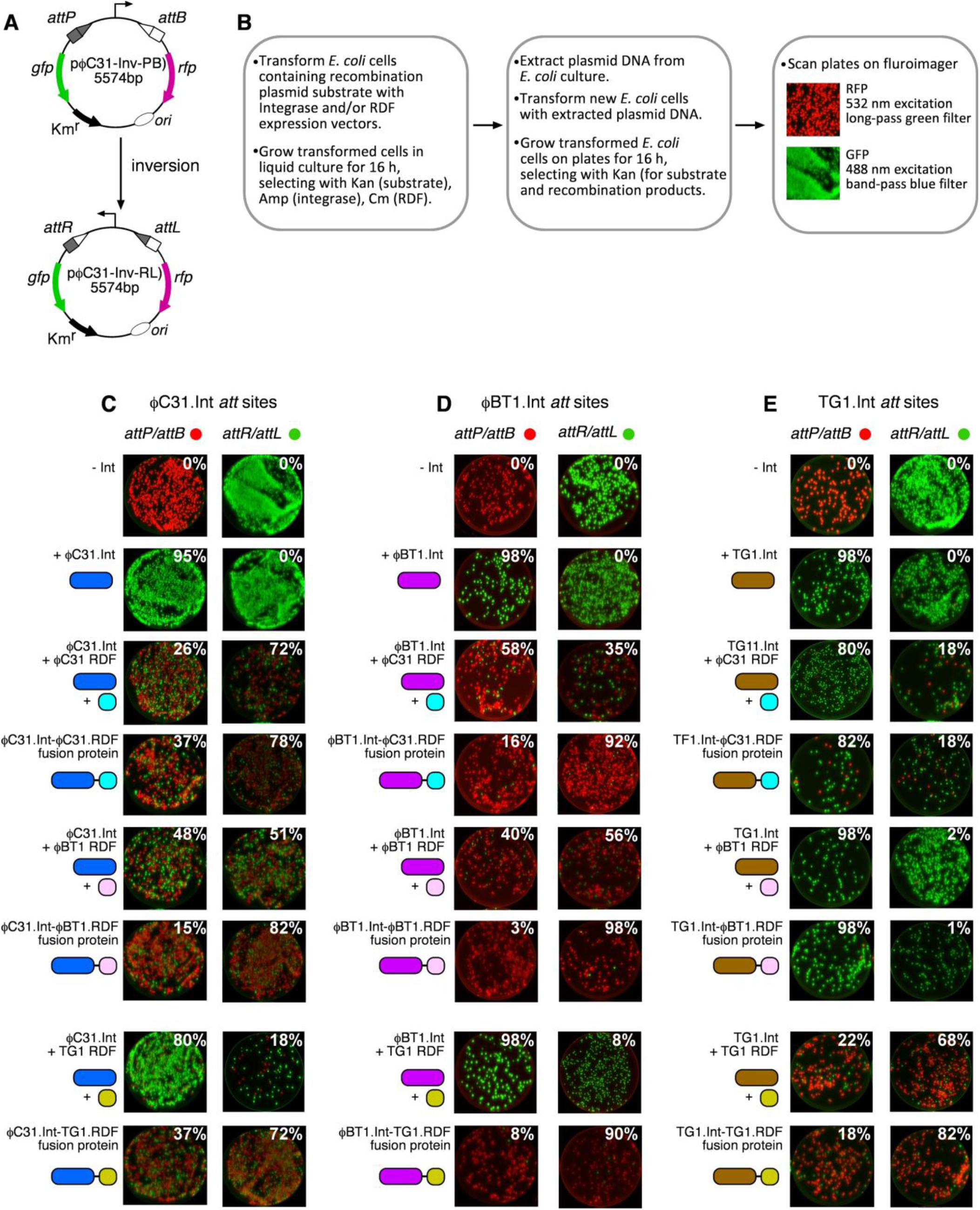
*In vivo* recombination reactions of ϕC31, ϕBT1, and TG1 integrases. (**A**) Scheme illustrating the *in vivo* intramolecular recombination (inversion) assay. In its default state, the promoter constitutively drives the expression of a red fluorescent protein (*rfp*) gene (pink arrow). A terminator sequence upstream of the promoter inhibits transcriptional read-through to the green fluorescent protein (*gfp*) gene (green arrow). Upon integrase-catalysed site-specific inversion reaction, the orientation of the promoter is flipped to allow the expression of GFP, and block RFP production. (**B**) Summary of the protocol for in vivo recombination reactions using constitutive integrase and RDF expression vectors. (**C**) Recombination activities of ϕC31 integrase in the presence of ϕC31-RDF, ϕBT1-RDF, and TG1-RDF; and as integrase-RDF fusions. In *attP*/*attB* reactions, cells start expressing RFP and produce GFP upon recombination. The extent of recombination is indicated as percentage of cells expressing GFP as outlined in **B** above. The reverse applies in reactions where the starting substrates are attR x attL. (**D**) Recombination activities of ϕBT1 integrase in the presence of ϕC31-RDF, ϕBT1-RDF, and TG1-RDF; and as integrase-RDF fusions. (**E**) Recombination activities of ϕTG1 integrase in the presence of ϕC31-RDF, ϕBT1-RDF, and TG1-RDF; and as integrase-RDF fusions. In panels **C, D**, and **E**, integrases are depicted as long ovals, and RDFs as short ovals. Each integrase and its cognate RDF are colour-coded to highlight expected orthogonal interactions.

**Figure 6:**
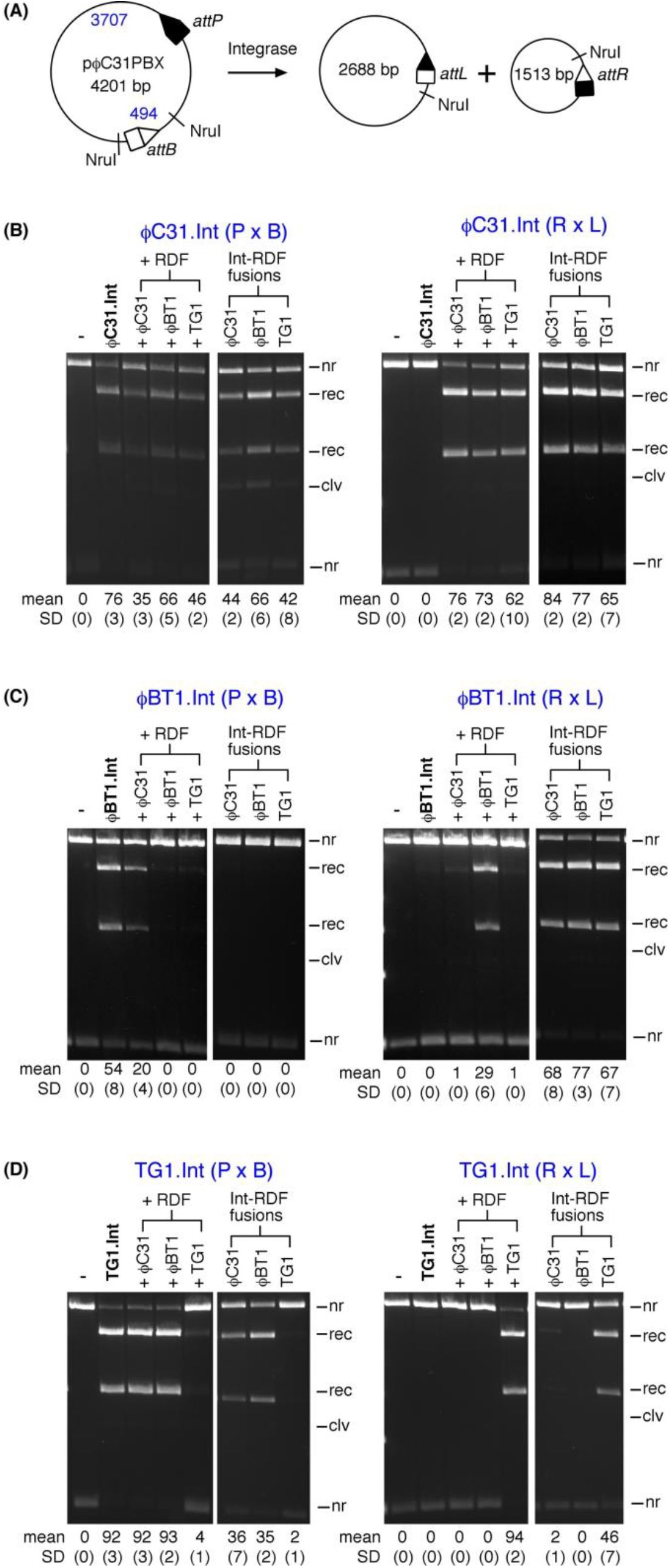
*In vitro* recombination reactions of ϕC31, ϕBT1, and TG1 integrases. (**A**) Scheme illustrating the *in vitro* intramolecular recombination assay (substrate plasmid pϕC31PBX for ϕC31 integrase is illustrated; the substrates for ϕBT1 and TG1 integrases are of the same design). The plasmid substrates are named after the LSI (ϕC31) and the *att* sites recombining (PBX; *attP* X *attB* resolution reaction). Upon recombination, the plasmid substrate gives two circular products in which the *attR* and *attL* sites are separated. For the reverse reaction, the starting substrate plasmid has *attP* and *attB* sites replaced by *attR* X *attL* sites, respectively, with recombination giving *attP* and *attB* sites on separate circular plasmid products. (**B**) Recombination activities of ϕC31 integrase in the presence of ϕC31-RDF, ϕBT1-RDF, and TG1-RDF. Reactions were incubated for 2 hours in the reaction buffer described in Materials and Methods. Reaction products were digested with the restriction endonuclease NruI prior to 1.2% agarose gel electrophoresis. In reactions where the integrase and RDF are added as separate proteins, the final concentration of both proteins were 200 nM. When the reactions were carried out using integrase-RDF fusion, the final concentration was 200 nM. The bands on the gel are labeled *nr* (non-recombinant, i.e. substrate), *rec* (recombination product). The mean extent of recombination and standard deviation (%) from quantitation of triplicate experiments are given below each lane. (**C**) Recombination activities of ϕBT1 integrase in the presence of ϕC31-RDF, ϕBT1-RDF, and TG1-RDF. Reaction conditions, gel electrophoresis, data acquisition and analyses areas described above in (A). (**D**) Recombination activities of TG1 integrase in the presence of ϕC31-RDF, ϕBT1-RDF, and TG1-RDF. Reaction conditions, gel electrophoresis, data acquisition and analyses are as described in (A).

However, *in vitro* recombination did not go to completion after 2 hours, with ϕC31, ϕBT1, and TG1 integrases converting 76%, 54%, and 92% of the substrate, respectively. We cannot differentiate from these data whether the lower amount of *in vitro* product for ϕC31 and ϕBT1 integrases can be ascribed to an intrinsically lower initial reaction rate or instability of the protein over 2 hours under the conditions used. The higher completeness of recombination observed *in vivo* could be due to the continuous expression of the proteins over the 16-hour growth period. The high activity of TG1 integrase is particularly noticeable, outperforming ϕC31 integrase, a recombinase that has been used in several *in vitro* and *in vivo* applications. Overall, the activities of the three integrases are consistent with our findings reported in an earlier study where we compared the activities of 10 different integrases (Abioye *et al*., 2022).

### Effects of RDFs on the recombination activities of ϕC31, ϕBT1, and TG1 integrases

To study the specificity of Integrase-RDF interactions across the three integrases, we studied the activities of each integrase in the presence of the three different RDFs both *in vivo* and *in vitro*. We did this in two ways: First, by using the integrase and the RDF as separate proteins, and secondly by constructing integrase-RDF fusions (Olorunniji *et al*., 2017) to account for effects due to differential binding affinities of integrases for non-cognate RDFs. Use of fusions also avoids potential differences in expression levels of the integrase and RDF proteins. In addition to activating *attR* x *attL* recombination, RDFs inhibit recombination of *attP* x *attB* reaction by their respective integrases (Khaleel *et al*., 2011; Bibb *et al*., 2005; Ghosh *et al*., 2006). To see if there is a correlation in activity levels between RDF-mediated activation of excisive recombination and inhibition of integrative recombination, we also investigated the inhibition of *attP* x *attB* recombination by the cognate RDF for the three integrases.

*In vivo*, the ϕBT1 RDF, when fused to ϕBT1 integrase (and to a lesser extent ϕC31 integrase), was the most effective in both activating excisive recombination and inhibiting integrative recombination (Figure 5). Furthermore, plotting the reaction endpoints for all of the pairwise tests shown in Figure 5 gives a strong anti-correlation between the endpoints of the *attP* x *attB* and the *attR* x *attL* reactions: the data plotted in Figure 7a have correlation coefficient of -0.99. This confirms that equilibrium was reached in these *in vivo* assays, and that the effectiveness of a particular RDF in promoting *attR* x *attL* reactions directly correlates with its effectiveness in inhibiting *attP* x *attB* reactions: each pair tested reached its particular “set point” regardless of the starting conditions.

**Figure 7:**
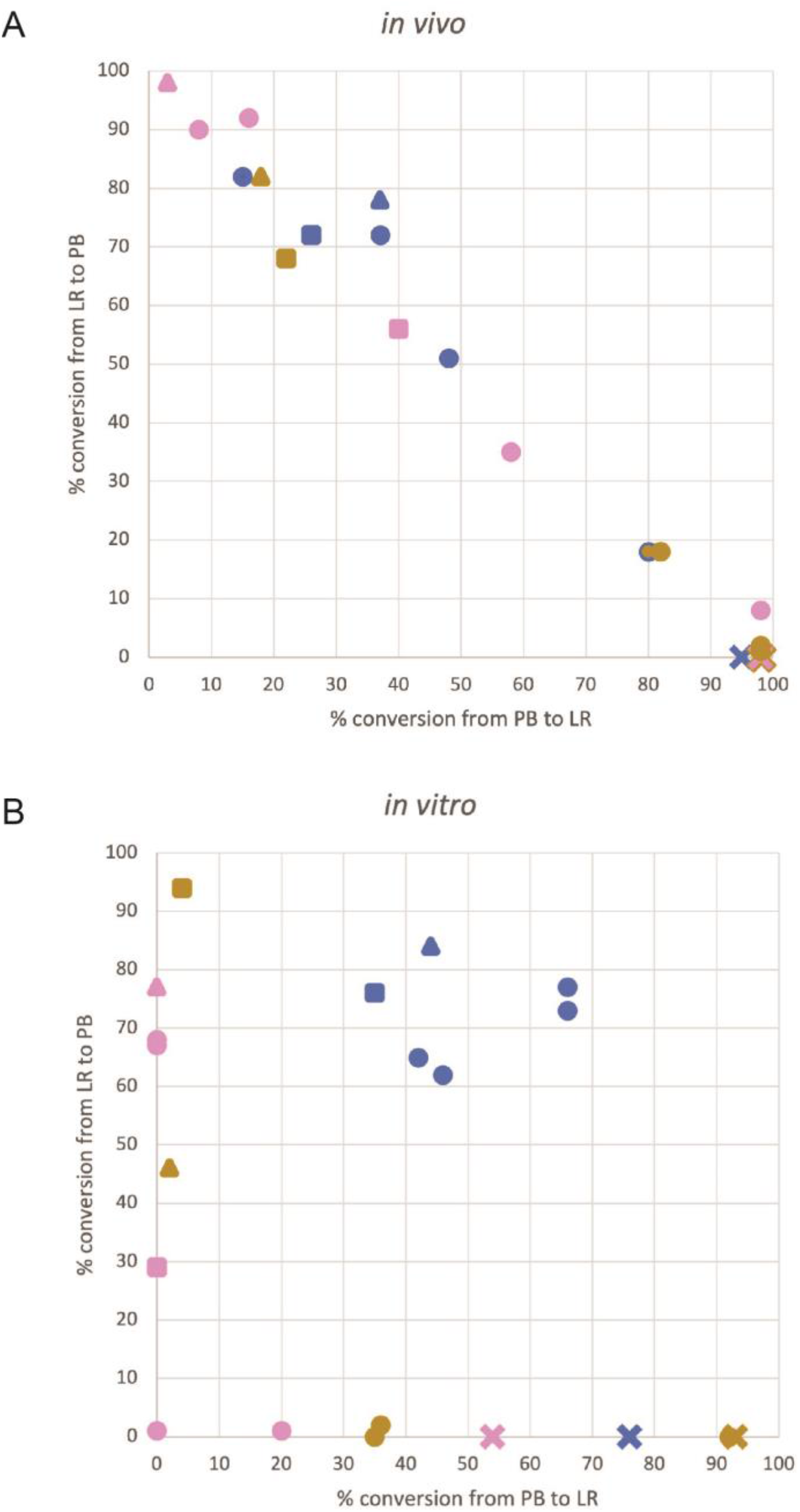
Correlation of extent of recombination in *attP* x *attB* and *attR* x *attL* reactions in the presence of the recombination directionality factor. (A) *In vivo* recombination, showing data taken from Figure 5. (B) *In vitro* recombination, showing data taken from Figure 6. The colour scheme is the same as in Figures 5 and 6: blue, ϕC31 integrase; pink, ϕBT1 integrase, gold, TG1 integrase. Triangles denote a cognate RDF fused to the integrase; square, a cognate RDF as a separate protein; circle, a non-cognate RDF; and X, no RDF.

*In vitro*, in the presence of their cognate RDFs, ϕC31, ϕBT1, and TG1 integrases recombined 76%, 29%, and 94% respectively of their *attR* x *attL* substrates. As for *attP* x *attB* recombination, TG1 integrase is the most active among the three, giving near complete conversion of the substrate plasmid (Figure 6). The overall pattern emerging from this shows that TG1 integrase is the most active among the three, giving near complete recombination in both integrative and excisive reactions. Incomplete reactions *in vitro* could be due to the factors discussed above for the *in vitro attP* x *attB* reactions, as well as weak directionality for the *attR* x *attL* reaction: that is, the equilibrium constant for the *attR* x *attL* reaction in the presence of RDF may not lie as far in favour of products vs. substrates as it does for the *attP* x *attB* reaction in the absence of RDF. This is supported in relation to the ϕC31 integrase by the conversion of 35 – 44% of the *attP* x *attB* substrate to product in the presence of the RDF. In contrast, the *in vitro* data for the ϕBT1 integrase in the presence of its RDF suggests that it simply did not reach equilibrium under the conditions used.

Plotting the reaction extent for each *in vitro* experiment (Figure 7b) highlights additional aspects of these reactions. Unlike the *in vivo* inversion assays, the *in vitro* assays monitor deletion of a plasmid segment. Therefore, the expected equilibrium of a given reaction is more complicated to predict. While the forward reaction depends on intramolecular synapsis of two *att* sites within the same plasmid, the reverse reaction requires intermolecular synapsis between *att* sites on separate DNA circles that may have diffused away from one another. If formation of the intermolecular synapse is too difficult under the conditions used, the endpoints would be expected to lie on the axes, as they do for ϕBT1 and TG1. In contrast, if the barrier to intermolecular synapsis is not significantly different from the barrier to intramolecular synapsis, the reaction endpoints would be expected to lie on the diagonal, similar to what is seen in Figure 7a. That is indeed approximately the case for ϕC31, indicating that ϕC31 integrase may form intermolecular synaptic complexes more readily than ϕBT1 and TG1 integrases do.

*In vitro*, ϕBT1 and TG1 RDFs were noticeably effective at inhibiting *attP* x *attB* recombination by their respective integrases, giving less than 5% activity in both cases. Complete inhibition of *attP* x *attB* reaction by the RDF is a key feature necessary for use of Integrase/RDF pairs in applications where integrases are used as binary genetic switches. TG1 integrase will be particularly suitable for building such devices since it shows near complete integrative and excisive activities. Figure 7a also shows that different integrase – RDF pairs could be used in applications where a tunable switch is required (e.g. a promoter inversion reaction that is partially rather than fully biased toward one outcome).

### The effect of covalent integrase-RDF linkage

We used integrase-RDF fusions (Olorunniji *et al*., 2017) to further investigate the specificity of integrase/RDF interactions across the three integrases and their RDFs. Among the three integrases, ϕC31 integrase showed the least orthogonal behaviour, responding to *attR* x *attL* activation and *attP* x *attB* inhibition by all three RDFs in both *in vivo* and *in vitro* reactions. In all cases, ϕC31-RDF was not as effective at regulating the activities of the integrase (Figures 5 and 6).

As expected, ϕBT1 integrase prefers its cognate RDF in regulation of *attR* x *attL* and *attP* x *attB* recombination (Figure 6). However, there is a noticeable difference in the interaction of ϕBT1 integrase with the RDFs when the two proteins are supplied separately and when they are fused together. ϕBT1 integrase had limited affinity for ϕC31-RDF and TG1-RDF when the RDFs were used as separate proteins. However, when the non-cognate RDFs were fused to ϕBT1 integrase, they were more effective in activation of *attR* x *attL* recombination and inhibition of *attP* x *attB* recombination (Figure 6).

In contrast to ϕC31 integrase and ϕBT1 integrase, TG1 integrase showed a high degree of selectivity for its cognate RDF. TG1 integrase prefers its cognate RDF and showed insignificant activity toward ϕC31-RDF and ϕBT1-RDF, either supplied as separate proteins or when fused to the integrase. This is especially noticeable in *in vitro* reactions, the exception being ϕC31-RDF showing *attR* x *attL* activation when fused to TG1 integrase (Figure 6).

## Discussion

### *In vivo* and *in vitro attP* x *attB* activities of ϕC31, ϕBT1 and TG1 integrases

Activities of these three integrases have been reported in a previous *in vitro* study in which the properties of 10 integrases were compared (Abioye *et al*., 2022). Several earlier reports have observed that Bxb1 integrase is the most active integrases characterised *in vitro* (Russell *et al*., 2006; Xu *et al*., 2013; Duportet *et al*., 2014; Jusiak *et al*., 2019).

Among the three integrases studied here, TG1 Integrase is the most active in recombining its *attP* and *attB* sites. Despite the high degree of sequence similarity, the recombination activity of TG1 integrase is noticeably higher than that of the other two integrases. Since there are no obvious differences in the sequences of the integrases around the catalytic residues, it is likely that the observed differences are due to other factors, including the fit of the att sites for each integrase – while evolutionary pressure on the phage may select an optimal *attP* sequence for integrase action to attain lysogeny, evolutionary pressure on the bacterial host may have the opposite effect on the sequence of *attB* (Ghosh *et al*., 2005; Mandali *et al*., 2013; McEwan *et al*., 2009; Adams *et al*., 2004). Presuming the fitness of each integrase for its natural biological roles, these differences in *E. coli* and *in vitro* might ‘accidentally’ reflect the ability of each integrase to adapt to unnatural conditions by integrating at non-cognate *attB* sites. It is also possible that the integrases have unknown factors in their natural contexts that stimulate their activity, but which are absent in these assays.

### Efficiency and specificity of RDF-dependent integrase activities

It is not surprising that there is a degree of cross-reactivity between the integrases and their RDFs. Among the three integrases, ϕC31 integrase was the least selective, with all three RDFs being able to activate it for *attR* x *attL* recombination. It is not clear why ϕC31 integrase is less selective than TG1 and ϕBT1 integrases, but these findings suggest a limitation in its use in the presence of the other two integrases.

In contrast to ϕC31 integrase, TGI integrase is highly selective and interacts with its RDF, gp25, to promote orthogonal clean switching reactions. For applications requiring control of directionality by the RDF, clean switching integrase-RDF pairs are essential. The findings here suggest that TG1 integrase would be suitable for such systems.

The affinity of ϕBT1 integrase for non-cognate RDFs (of TG1 and ϕC31) is significantly increased by covalent attachment of such RDFs. When used as separate proteins, ϕBT1 integrase did not interact significantly with these non-cognate RDFs. However, covalent attachment resulted in the ability of TG1 and ϕC31 RDFs to activate *attR* x *attL* recombination and inhibit *attP* x *attB* recombination by ϕBT1 integrase.

Overall, the observed weak integrase/RDF orthogonality among these three enzymes emphasizes the need for identifying more integrases with known RDFs. To date, only a handful of active Integrase/RDF have been characterised, *in vivo* and/or *in vitro*. These include Bxb1 (Ghosh *et al*., 2006), ϕRV1 (Bibb *et al*., 2005), ϕC31 (Khaleel *et al*., 2011), ϕBT1 (Zhang *et al*., 2013), TP901 (Breuner *et al*., 1999), A118 (Mandali *et al*., 2017), SPBc (Abe *et al*., 2017), ϕJoe (Fogg *et al*., 2017), CD1231 (Serrano *et al*., 2016). In addition, the RDF for TG1 integrase (gp25) was identified by Zhang *et al*., (2013), but its activity has not been demonstrated *in vivo* or *in vitro*.

The availability of a larger set of integrases with specific RDFs, with known clean switching activities and orthogonal to each other, will facilitate the use of serine integrases in designing genetic circuits and other regulated systems in synthetic biology. Additionally, the plot if Figure 7a shows that tunability could be added to existing genetic circuits by using related but not strictly cognate RDFs for a given integrase: in that way, the set point of a particular inversion switch could be changed simply by changing the RDF.

Our findings also provide an opportunity for identifying the protein-protein interactions among this family of integrase/RDF pairs that determine specificity for the integrase. For example, the strict specificity of TG1 integrase for its RDF in contrast to the broad tolerance of ϕC31 integrase to all three RDFs could be a starting point for identification of protein features and interactions that determine *attR* x *attL* activation, *attP* x *attB* inhibition, and selectivity for the cognate integrase.

### AlphaFold2 multimer predicts a model for integrase/RDF interaction

The predicted interactions of each of the three integrases with their cognate RDFs are very similar. However, the sequence conservation within the binding surface on the RDF is much more conserved than that on the integrase (Figure 8). This could be an indication that specificity or selectivity is determined more by the integrase than the RDF. The significant exception is seen in TG1 integrase, which has a small insertion that could allow it to make a more stable or more rigid interaction with the RDF (Figure 8). It is likely that this insertion could explain the enhanced ability of TG1 integrase to discriminate among RDFs (Figures 5 and 6). Investigating the effects of deleting this element from TG1 integrase or inserting it into ϕC31 or ϕBT1 integrases could provide some insights into the design and understanding of future genetic switches.

**Figure 8:**
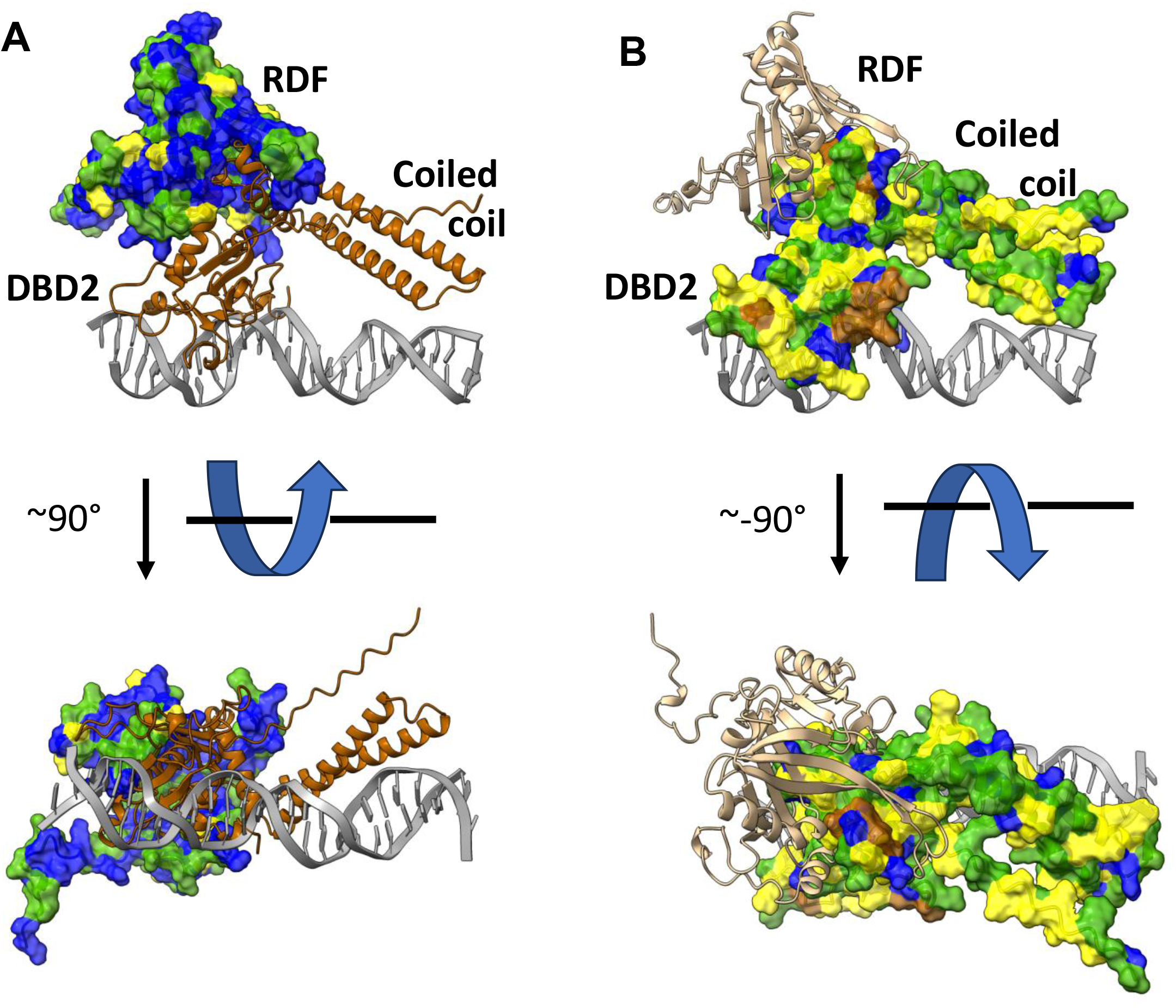
AlphaFold2-predicted structures of the second DNA-binding domains (DBD2) of ϕC31, ϕBT1, and TG1 integrases and their RDFs. These models were mapped domain-wise onto the individual domains of 4kis.pdb as described in Figure 4. TG1 Integrase is shown with conservation mapped to semi-transparent surface (on RDF on left; int on right). Yellow: none; green: identical in 2; blue: identical in all 3; brown: insertions in TG1 integrase relative to other two. Note that the binding surface on RDF is much more conserved than binding surface on Integrase.

## Materials and Methods

### *In vivo* recombination reactions

Plasmids for constitutive expression of integrases, RDFs, and integrase–RDF fusion proteins in *E. coli* were made as described in Olorunniji *et al*. (2017). Recombination activities (inversion) of ϕC31 integrase were carried out using the plasmid substrate, pϕC31-invPB and pϕC31-invRL, as described in Olorunniji *et al* (2019). Similar plasmid substrates for ϕBT1 and TG1 integrases were made by cloning the corresponding att sites into pϕC31-invPB or pϕC31-invRL. Recombination activity was measured using the invertible promoter reporter system (see Figure 5). To assay ϕC31 integrase *attP* x *attB* recombination, *E. coli* DS941 cells containing the pϕC31-invPB substrate were transformed with the vector plasmid expressing ϕC31integrase. The extent of *attP* x *attB* recombination was monitored by measuring the extent of switching from RFP to GFP expression. The extent of recombination was determined by counting the number of colonies expressing either RFP or GFP. TG1 and ϕBT1 integrases were assayed the same way using their corresponding *in vivo* substrates. Scanning of *E. coli* cell fluorescence was carried out using a Typhoon FLA 9500 fluorimager (GE Healthcare) as described in Olorunniji *et al*. (2019). Briefly, fluorescence of the expressed proteins was measured (GFP: excitation, 488 nm, band-pass blue filter; RFP: excitation, 532 nm, long-pass green filter).

### Expression and purification of serine integrases

The three integrases were expressed in E. coli BL21(DE3)pLysS and purified as described in Olorunniji *et al*. (2017). Briefly, the expression strain for each integrase was grown at 37 ° C in 2x YT-broth to A_600_ of 0.6 to 0.8. Cultures were cooled to 20 ° C and integrase expression was induced with 0.75 mM IPTG, after which the cultures were grown for 16 hours at 20 ° C. The proteins were purified by nickel affinity chromatography and bound proteins were eluted with an imidazole gradient buffer system. Fractions were collected and SDS-PAGE was used to determine peaks corresponding to the proteins of interest. Fractions containing the integrases were dialysed against Protein Dilution Buffer, PDB (25 mM Tris–HCl (pH 7.5), 1 mM DTT, 1 M NaCl and 50% glycerol), and stored at –20 ° C. Dilutions of each integrase for *in vitro* recombination reactions were made into the same buffer.

### *in vitro* recombination reactions

Substrate plasmids for the assay of intramolecular activities of the three integrases *in vitro* are described in Abioye *et al*. (2022). The plasmid substrate for each integrase is named according to the integrase *att* sites. Figure 6 shows the substrate for ϕC31 integrase in which pϕC31PBX carries ϕC31 *attP* and *attB* sites. The *att* sites are arranged in a ‘head to tail’ orientation leading to resolution of the substrate plasmid into two smaller product plasmids upon recombination. *In vitro* recombination of supercoiled plasmid substrates and analysis of recombination products were carried out as reported in Abioye *et al*. (2022). Typically, recombination reactions were carried out by adding integrase (2 µM, 5 µl) to a 30 µl solution containing the plasmid substrate (25 µg/ml), 50 mM Tris-HCl (pH 7.5), 100 µg/ml BSA, 5 mM spermidine, and 0.1 mM EDTA. Samples were incubated at 30 ° C for 2 hours, after which the reactions were stopped by heating at 80 ° C for 10 minutes. The samples were cooled and treated with NruI (New England Biolabs) to facilitate analysis of recombination products. Following the digest, samples were treated with SDS and protease K before reaction products were separated by agarose gel electrophoresis (Abioye *et al*., 2022).

### AlphaFold2-based protein structure prediction

Three-dimensional (3D) protein structures were generated using the colabfold implementation AlphaFold2-multimer; version 1.5.2 with default parameters (Miridita *et al*., 2022; Jumper *et al*., 2021; Evans *et al*., 2022). The structures were viewed and manipulated using PyMol (https://pymol.org) (Figure 4) and UCSF ChimeraX (Goddard *et al*., 2018) (Figure 7). The models shown were predicted using full-length integrase and RDF sequences. Nearly identical interactions between DBD2 and the RDF were predicted for each pair when the integrase sequences were truncated to include only DBD2 and the coiled coil.

## Acknowledgments

Molecular graphics and analyses were performed with UCSF ChimeraX, developed by the Resource for Biocomputing, Visualization, and Informatics at the University of California, San Francisco, with support from National Institutes of Health R01-GM129325 and the Office of Cyber Infrastructure and Computational Biology, National Institute of Allergy and Infectious Diseases.

